# Different mechanisms for modulation of the initiation and steady-state of smooth pursuit eye movements

**DOI:** 10.1101/846030

**Authors:** Stuart Behling, Stephen G. Lisberger

**Author notes:** Proofs and correspondence to: Stuart Behling, Department of Neurobiology, Box 3209, Duke University School of Medicine, Durham, NC, 27710, Fax: obsolete.

## Abstract

Smooth pursuit eye movements are used by primates to track moving objects. They are initiated by sensory estimates of target speed represented in the middle temporal (MT) area of extrastriate visual cortex and then supported by motor feedback to maintain steady-state eye speed at target speed. Here, we show that reducing the coherence in a patch of dots for a tracking target degrades the eye speed both at the initiation of pursuit and during steady-state tracking, when eye speed reaches an asymptote well below target speed. The deficits are quantitatively different between the motor-supported steady-state of pursuit and the sensory-driven initiation of pursuit, suggesting separate mechanisms. The deficit in visually-guided pursuit initiation could not explain the deficit in steady-state tracking. Pulses of target speed during steady-state tracking revealed lower sensitivities to image motion across the retina for lower values of dot coherence. However, sensitivity was not zero, implying that visual motion should still be driving eye velocity towards target velocity. When we changed dot coherence from 100% to lower values during accurate steady-state pursuit, we observed larger eye decelerations for lower coherences, as expected if motor feedback was reduced in gain. A simple pursuit model accounts for our data based on separate modulation of the strength of visual-motor transmission and motor feedback. We suggest that reduced dot coherence creates less reliable target motion that impacts pursuit initiation by changing the gain of visual-motor transmission and perturbs steady-state tracking by modulation of the motor corollary discharges that comprise eye velocity memory.

## Introduction

A critical component of cerebellar organization and function is integration of sensory information about the environment with copies of motor commands to support sensory-motor behavior (Huang et al., 2013). Smooth pursuit eye movements are an example motor behavior where the cerebellum integrates sensory and motor signals (Stone and Lisberger, 1990; Churchland and Lisberger, 2000, 2001; Lisberger, 2009). Pursuit is used by primates to track moving targets. Visual motion pathways drive the initial eye response to initiate pursuit. By contrast, visual motion signals play a minimal role during steady-state tracking (Lisberger et al., 1987), when previous eye movement commands sustain eye velocity by feedback of an “extraretinal signal” (Morris and Lisberger, 1987; Churchland and Lisberger, 2001; Churchland et al., 2003; Madelain and Krauzlis, 2003), at least to some degree through the cerebellum (Miles and Fuller, 1976; Lisberger and Fuchs, 1978; Stone and Lisberger, 1990; Shidara et al, 1993).

A representation of target speed in the middle temporal (MT) area of extrastriate visual cortex drives the initiation of pursuit (Newsome et al., 1985; Priebe et al., 2003). The pursuit system uses the initial response of the population of neurons in MT to estimate target speed and direction. However, the eye speed at the initiation of pursuit depends not only on the representation of target motion in MT, but also on the sensed reliability of the motion. Previous work from our lab has demonstrated that some target forms provide a less reliable representation of motion and that lower motion reliability leads to lower eye speed during pursuit initiation (Darlington et al., 2017). We think of this modulation in terms of control of the strength or “gain” of visual-motor transmission, and several lines of evidence implicate the smooth eye movement region of the frontal eye fields in both gain control and the assessment of motion reliability (Tanaka and Lisberger, 2002; Nuding et al., 2009; Darlington et al., 2017).

Visual motion cannot sustain steady-state tracking once eye speed matches target speed because retinal image motion is effectively zero and motion sensitive neurons in area MT would be unresponsive because they respond selectively in the presence of image motion (Maunsell and Van Essen, 1983). Instead, eye velocity seems to be sustained by a neural signal that is related more to target velocity than to image velocity (Yasui and Young, 1975; Robinson et al., 1986). Recordings from the floccular complex of the cerebellum suggest that positive feedback based on corollary discharges of the motor command could provide a key mechanism for implementing a neural correlate of target velocity (Miles and Fuller, 1976; Lisberger and Fuchs, 1978; Stone and Lisberger, 1990). Floccular Purkinje cells receive input from the sensory pathway as well as positive motor feedback and generate simple-spike firing that represents a kinematic model of eye movements (Shidara et al., 1993; Medina and Lisberger, 2009). In turn, Purkinje cells have disynaptic connections to the motoneurons for the eye muscles and therefore drive pursuit (Highstein, 1973; Scudder and Fuchs, 1992; Lisberger et al., 1994).

Most previous thinking about the role of the floccular complex has assumed that eye velocity positive feedback is fixed and immutable (Stone and Lisberger, 1990; Krauzlis and Lisberger, 1994; Schwartz and Lisberger, 1994). However, our analysis of the expression of motor learning in pursuit raised the possibility that it is subject to modulation (Yang and Lisberger, 2010; Hall et al., 2018). To determine if the steady-state of pursuit is modulated by motion reliability, and if the modulation is separate from that for the initiation of pursuit, we have developed a target and a task that reliably perturbs both pursuit initiation and steady-state tracking. We use motion coherence in a patch of dots (Newsome and Paré, 1988) as a way to change target form and motion reliability without changing the speed or direction of target motion. We observed that pursuit initiation and steady-state tracking responded in quantitatively different ways to changes in dot coherence. Computational modeling demonstrates how separate modulation of the strength of visual-motor transmission and eye velocity positive feedback can reproduce the relationships observed in the data.

## Methods

The two adult male rhesus monkeys used in these experiments weighed between 12 and 16 kg. All experimental protocols were approved by the *Institutional Animal Care and Use Committee* at Duke University and all procedures were in accordance with the National Institutes of Health *Guide for the Care and Use of Laboratory Animals*. A scleral coil to track eye position and a headpost to restrain their heads during experiments were surgically implanted (Ramachandran and Lisberger, 2005) using sterile procedure while each monkey was under general anesthesia with isofluorane. Analgesics were given postsurgically. Monkeys then were trained to fixate and pursue moving targets in exchange for juice rewards, using the analog signals from the eye coil to determine the movements of the eye. Both monkeys had substantial pursuit experience before these experiments were carried out.

### Visual Stimuli and Behavioral Task

Visual targets were presented on a 23” CRT monitor with a refresh rate of 80 Hz. The monitor was 30 cm from the monkey’s eyes for a field of view of 77 by 55 degrees. The background was a neutral grey in relation to the stimulus colors of white and black.

Experiments consisted of a series of brief eye movement behavioral trials. Trial structure and target form was created and controlled using ‘Maestro’, a software system created by our laboratory. The trials used a single black circular dot (0.6 x 0.6 degrees) for fixation spots and patches of dots as pursuit targets (Yang et al., 2012). The individual dots were 5 pixels in diameter and patches of dots contained 72 dots within a 4-degree diameter invisible aperture. Half of the dots were bright and half dark so as not to change the average luminance when the target was presented. We varied the motion coherence of the dot patch by changing the probability that each dot would maintain the trajectory from previous frames versus undergoing random replacement within the aperture (Figure 1a). Lower coherence targets involved higher probabilities that any given dot would be replaced at a random location, resulting in greater amounts of motion noise that masks the motion in the stimulus (Britten et al., 1996; Schütz et al., 2010).

**Figure 1.**
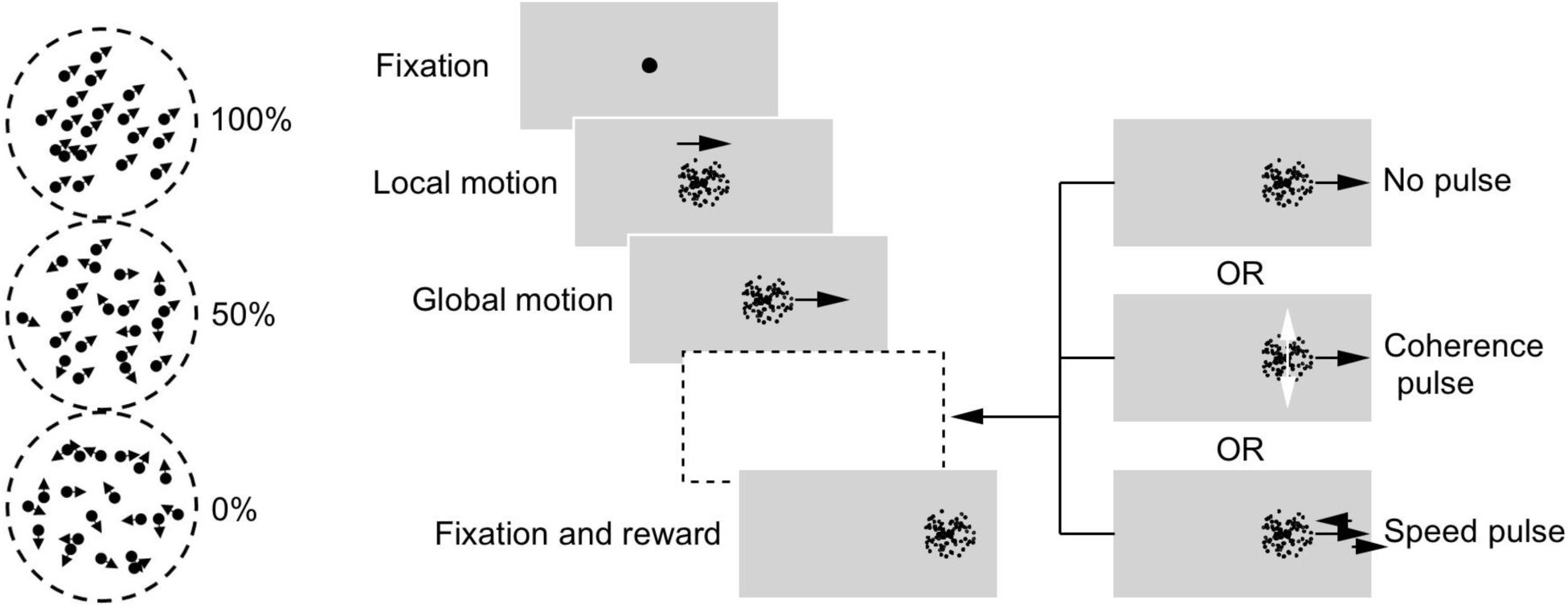
Stimulus and behavioral paradigm. The three patches of dots on the left show the motion of dots within patches that have 100%, 50%, or 0% coherence. The schematic on the right shows the trial structure. Each trial begins with fixation before moving through brief local motion of the dots within an invisible stationary aperture and global, *en bloc* motion of the dots and the aperture to a fixation stationary stimulus for fixation and reward. The same dot coherence was retained during the periods of both local and global motion. After the start of global motion, the target can continue unaltered (no pulse), or can undergo either a brief pulse of either dot coherence or stimulus speed.

Trials (Figure 1b) always started with fixation for a randomly selected duration between 600 and 1000 ms. During fixation, the trials were aborted if the monkey’s eyes strayed more than 2 degrees from the target. After successful fixation, the fixation target disappeared and was replaced with a patch of moving dots. Each trial started with local motion of dots within a stationary but invisible aperture for 100 ms to reduce the probability of early catch-up saccades (Osborne et al., 2007). Then, the local motion transitioned seamlessly to global motion. During global motion, we retained the dot coherence used during location motion, and also moved the invisible aperture at the speed selected for the trial. Global motion continued for a random interval from 900 to 1500 ms before the trial ended with fixation of a stationary spot for 500 ms to ensure completion of the entire trial. In select trials, stimulus parameters were perturbed 400 or 500 ms after the start of local motion with “speed pulses” or “coherence pulses” that had durations of 300 ms. For speed pulses, the speed of stimulus motion was increased or decreased by 1, 2, 4, or 8 deg/s. For coherence pulses, the dot coherence was raised or lowered while the stimulus speed remained unchanged. Targets with different movement parameters, dot coherences, and speed or coherence pulses were interleaved randomly. Trials were aborted if the monkey’s eyes strayed more than 4 degrees from stimulus position during the pursuit phase of the trial. Upon successful completion of a trial, the monkey received a juice reward through a sipper tube.

### Data Analysis

Filtered voltage signals proportional to horizontal and vertical eye velocity were obtained by feeding horizontal and vertical eye position voltage signals from the eye coil into an analog circuit that differentiated frequency content below 25 Hz and filtered out higher frequency content (−20 dB per decade above 25 Hz). For analysis, each trial was aligned to the start of local target motion and trimmed to the longest common trial length among all repetitions of a given stimulus. Saccades during pursuit were determined using acceleration and velocity thresholds of 1000 deg/s/s and 25 deg/s, respectively. Position and velocity values were treated as missing during times of saccades along with a buffer of 20 ms on each end of the saccade. If the automatic algorithm detected saccades during the first 200 ms of target motion, they were confirmed by visual inspection of the trace. Eye speed was calculated as a function of time from the horizontal and vertical eye velocity traces and then averaged at each millisecond across multiple repetitions of each target motion and form. Any period of the average eye speed trace that had saccades in over half of the trials was excluded from the records in our figures and from any additional analysis.

The onset of pursuit was determined as the time of maximum eye jerk (first derivative of acceleration) during the open-loop interval of pursuit initiation. The end of the open-loop interval is 100 ms after pursuit onset (Lisberger and Westbrook, 1985), when the first visual feedback is available about the results of initiating pursuit and the system starts to transition to steady-state tracking. To quantify the response of the eye during the open-loop interval and during steady-state tracking, we averaged eye speed in windows from 0 to 75 ms and 400 to 475 ms after pursuit onset, respectively. For analysis of the responses to speed or coherence pulses, peaks in the eye jerk were used to demarcate when the eye started to respond to the change in stimulus. For both speed and coherence pulses, we used the average eye acceleration over the first 75 ms of the response to measure the response to the pulse. Acceleration values were corrected by subtracting the same measurement using the same analysis window on data obtained at the identical target coherence but without a speed or coherence pulse.

## Results

One important feature of pursuit initiation is the delay of performance feedback. The time it takes the change in image motion caused by the initiation of pursuit to be processed is about 100 ms from the time of eye motion onset (Lisberger and Westbrook, 1985). We refer to this epoch of pursuit initiation as the open-loop interval and we use it as a window into the sensory processing driving pursuit. In contrast, steady-state tracking could use performance feedback but does not because retinal image motion is negligible when tracking is perfect or almost perfect. Instead, steady-state tracking seems to be supported by “eye velocity memory” that results from positive feedback of a corollary discharge of motor commands (Lisberger, 2009). Because of the different mechanisms involved in initiation versus steady-state pursuit, we study the two epochs of the overall movement separately. We use average eye speeds in the intervals from 0 to 75 ms and 400 and 475 ms after pursuit initiation to quantify pursuit initiation and steady-state tracking, respectively.

### Effects of dot coherence on pursuit initiation and steady-state

Across the initiation and steady-state of pursuit we saw graded decreases in eye speed as we reduced dot coherence. The data in Figure 2 show the time course of average eye speed from one day of pursing targets at 10 degree/s for each of two monkeys. We plot the same data in the left and right columns using an expanded time base to show the details of the responses during pursuit initiation (Figure 2A, C) and a contracted time-base to show the full responses, including steady-state tracking (Figure 2B, D). Eye speeds declined as a function of dot coherence as we reduced dot coherence in 10% decrements from 100% to 0%, during both pursuit initiation (0 to 75 ms after pursuit onset) and steady-state tracking (400 to 475 ms after pursuit onset) in both monkeys (Figure 2E, F).

**Figure 2.**
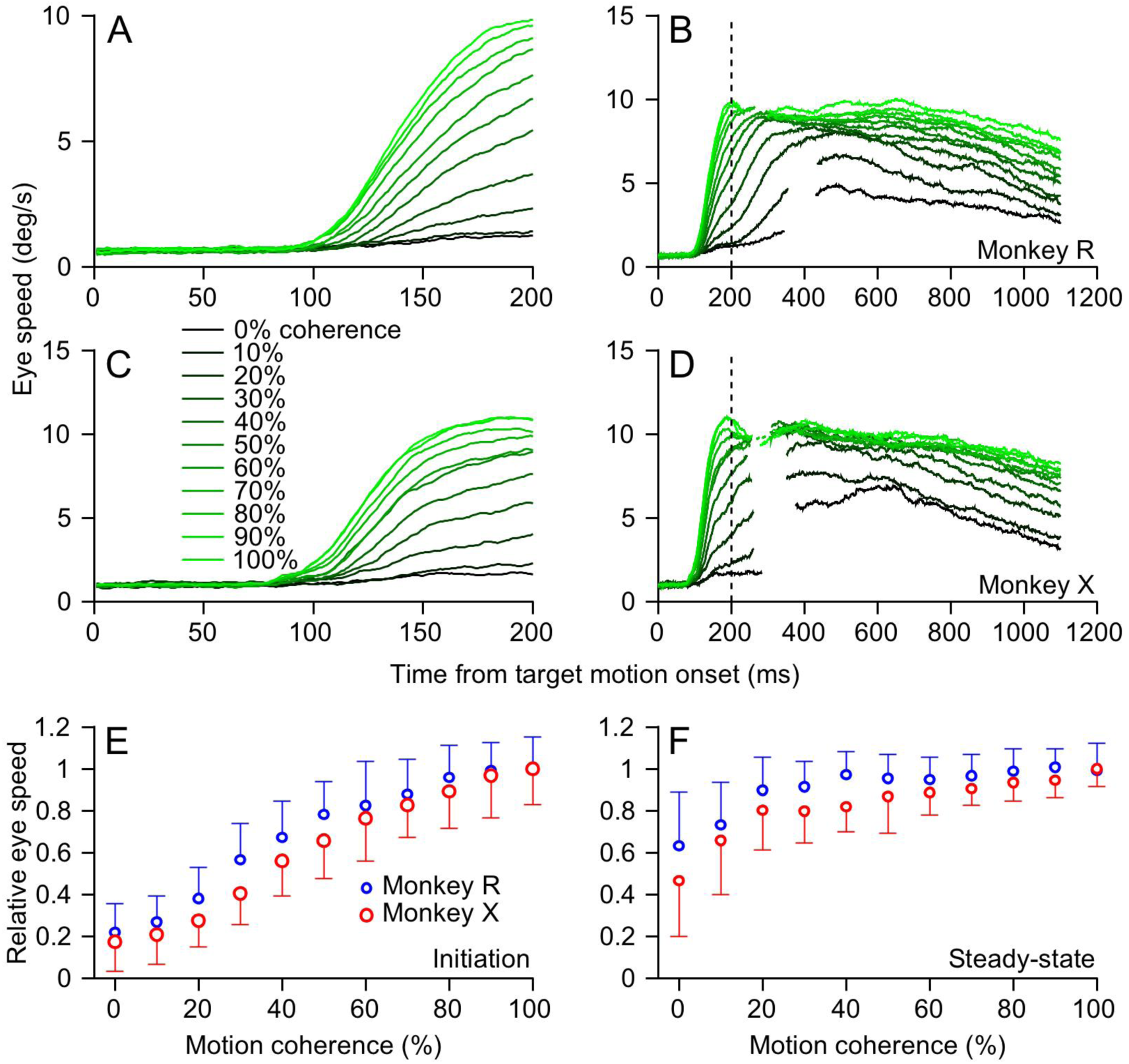
Effect of dot coherence on the initiation of pursuit and steady-state tracking. Average eye speed as a function of time for an example session in each of the two monkeys. **A, C**: Data for the first 200 ms after the onset of target motion at 10 deg/s. **B, D**: Data for the entire response to target motion, extending the example eye speed traces in **A** and **C**. The vertical dashed line marks 200 ms after the onset of target motion, where the traces in **A** and **C** end. The gaps in the eye speed traces indicate intervals when over half of trials used in the averages contained saccades. In **A-D**, the transition of colors from bright green to black indicates data for patches of dots moving with 100% to 0% coherence. **E, F**: Relative eye speed of data in A-D is plotted as a function of motion coherence. Error bars represent standard deviation across individual responses for a representative experiment in each monkey.

The effect of dot coherence on eye speed was quantitatively different for pursuit initiation and steady-state tracking. During initiation, both monkeys showed steady, almost linear decreases in relative eye speed as a function of dot coherence (Figure 3C, target speed 10 deg/s). During steady-state tracking, in contrast, relative eye speed decreased in a distinctively nonlinear way as a function of dot coherence (Figure 3A). To visualize the differences between these epochs, we plotted the relative effect on eye speed for pursuit initiation as a function of that effect on eye speed for pursuit steady-state for the same parameters of target motion and coherence (Figure 3B). The relationship is distinctly nonlinear. All points plot above the line of slope one, showing that coherence has larger effects on the initiation versus steady-state of pursuit over much of the range of coherence. The same relationship holds for target speeds of 20 deg/s. The differential impact of dot coherence on steady-state tracking versus pursuit initiation supports the idea of separate modulation of the signals that drive the two phases of pursuit.

**Figure 3.**
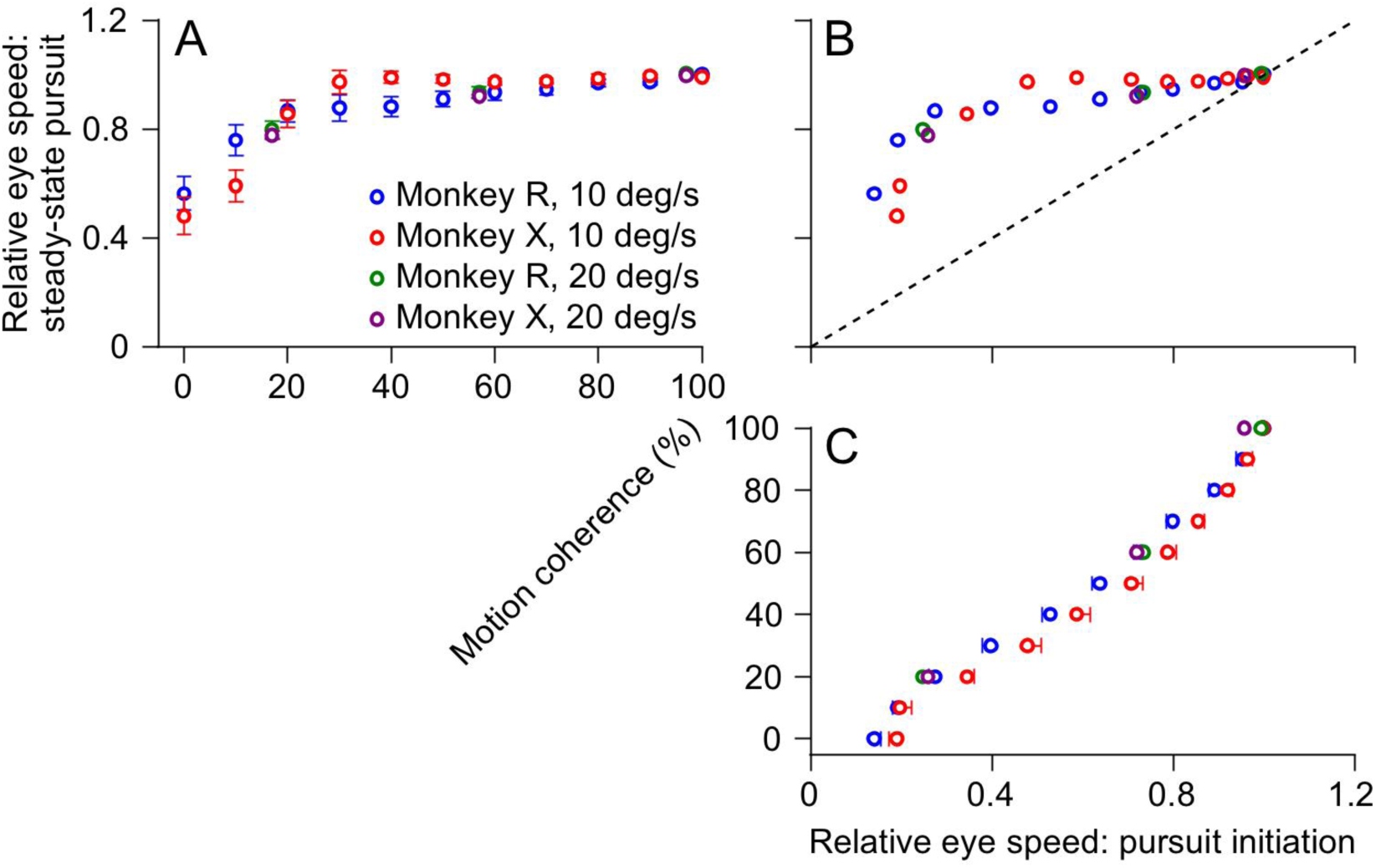
Quantitative comparison of the effects of dot coherence on the initiation versus the steady-state of pursuit. **A, C**: Relative eye speed is plotted as a function of motion coherence (or *vice versa* in **C**) for steady-state tracking (**A**) and the initiation of pursuit (**C**). **B**: Relative eye speed during steady-state tracking is plotted as a function of that during the initiation of pursuit for different values of dot coherence. The dashed line has a slope of one. In all 3 graphs, different colors show data for the two monkeys at two speeds. Data were normalized to have a value of 1 for motion coherence of 100% in each session and then averaged across sessions. Error bars are standard error of the mean across for n=4 experiments (Monkey R, 10 deg/s), n=6 experiments (Monkey X, 10 deg/s), n=10 experiments (Monkey R, 20 deg/s), and n=10 experiments (Monkey X, 20 deg/s).

Previous research has shown that decreases in the contrast of moving targets decrease eye speeds during pursuit initiation because of a reliability-weighted competition between sensory evidence and a prior that targets move slowly or not at all, where weaker or less reliable evidence decreases the gain of visuo-motor transmission (Darlington et al., 2018). We asked whether the same mechanism caused the effect of dot coherence on pursuit through an experimental design similar to that used in the prior studies (Darlington et al., 2017, 2018; Devaret et al., 2018). We measured the impact of priors created by blocks of mostly slow versus mostly fast target motions as we measured the pursuit evoked by high versus low coherence targets. We created priors through fast- or slow-context blocks that comprised 40 trials with target motion at 20 degree/s or 2 degree/s and 10 randomly-interleaved probe trials with target motion at 10 degree/s (right column of Figure 4). The 10 probe trials included 5 at each of 100% and 10% dot coherence, allowing comparisons to eye speed during control blocks that presented only probe trials with these two values of dot coherence.

**Figure 4.**
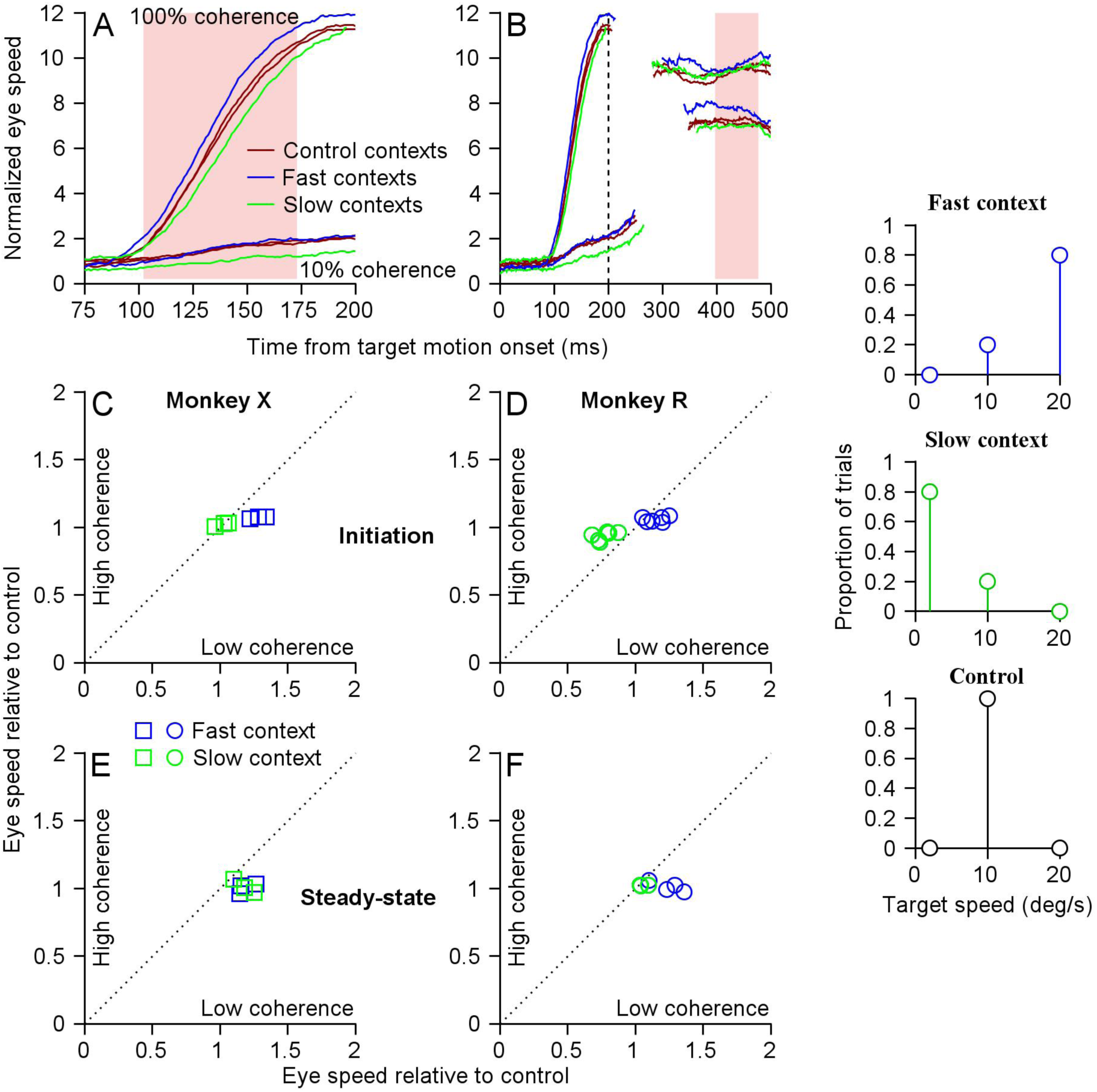
Effect of changing the prior for target speed on the eye speeds for low versus high coherence patches of dots. The three graphs in the right column show the mix of target speeds in each of the contexts used to control the speed prior. **A, B**: Averages of eye speed as a function of time from an example experiment in one monkey showing the initiation of pursuit (**A**) and steady-state tracking (**B**) for 100% and 10% coherence motion. Different colors show data for different speed contexts. The pink shading highlights the interval when eye speeds were analyzed. Gaps in the eye speed traces show the intervals when over half of the trials had saccades. **C-F**: Relative eye speeds for high coherence versus low coherence dots during pursuit initiation (**C, D**) and steady-state tracking (**E, F**). Dotted lines have slopes of one. Green and blue symbols show data for the slow and fast context, respectively. Each experimental day contributed a pair of symbols showing averages across at least 10 full cycles of slow, fast, and control contexts. All data are normalized for eye speeds during the control contexts.

As expected for a reliability-weighted impact of sensory information, we saw larger effects of context on the responses to lower coherence targets for pursuit initiation but only weak effects on steady-state pursuit (Figure 4). The expanded time base records in Figure 4A show faster eye speeds at the initiation of pursuit for the fast versus slow contexts (blue versus green traces), with larger percentage effects for low coherence targets. The longer time base records in Figure 4B show much smaller and less consistent effects on steady-state eye speed. To quantify the initiation and steady-state of pursuit separately, we averaged eye speeds from 0 to 75 ms and 300 to 375 ms after pursuit onset, respectively (light pink shading in Figure 4A, B). Trials were only 500 ms in duration for these experiments so we used an earlier interval to quantify steady-state pursuit. To avoid any single trial adaptation that might counteract the effect of context, we analyzed only probe trials that followed a trial at the context speed in the context blocks and did not analyze the first trial of control blocks following speed context blocks.

During pursuit initiation, context had a significant impact on eye speed for monkey X and R at high coherence (paired-sample t-test p=0.0048 and p<0.0001) and low coherence (paired-sample t-test p=0.0054 and p=0.0010). Regression of eye speeds at high and low coherence across speed contexts also demonstrated that the line of slope one lay outside the 95% confidence intervals for both monkeys during pursuit initiation. An N-way ANOVA supported a main effect of context on eye speeds for pursuit initiation in both monkeys (p<0.0001 in each). Therefore, context has stronger effects on pursuit eye speeds at low coherence than at high coherence during pursuit initiation (Figure 4C, D). Because context has a larger effect on the initiation of pursuit for low versus high coherence targets, Figure 4 supports the idea that deficits in eye speed from dot coherence result from a reliability-weighted competition between visual motion signals and previous experience (Darlington et al., 2017, 2018), rather than from misrepresentations of target speed. The data suggest that dot coherence, like contrast, affects the initiation of pursuit by changing the reliability of visual motion signals leading to reductions in visual-motor gain.

Eye speeds relative to control during steady-state tracking did not depend significantly on context for either high or low coherence dots (paired-sample t-test, p=0.0629 and p=0.1201 for monkey X and p=0.4007 and p=0.4157 for monkey R respectively). Because context did not consistently affect steady-state pursuit eye speeds, even for low coherence patches of dots, we cannot conclude that dot coherence modulates steady-state eye speed through a reliability-weighted competition between visual motion signals and previous experience. This provides additional evidence for separate mechanisms for modulation of the initiation versus steady-state of pursuit.

### Mechanisms of effects of dot coherence on pursuit

We begin our analysis of mechanism by verifying that the pursuit system remains engaged and operates in a machine-like way during steady-state tracking at low and high values of dot coherence. We document the machine-like behavior by measuring the eye speed response to momentary changes in (1) target speed or (2) dot coherence. For example, when we introduced changes in target speed, or speed pulses, during steady-state pursuit by either increasing or decreasing target speed by several deg/s for 300 ms (Figures 5A, B), we observed robust eye speed responses to speed pulses. The responses quickly converged towards the eye speeds seen in trials that used the same speed as each speed pulse throughout the trial for 100% coherence (Figure 5A) and 10% coherence (Figure 5B). Therefore, the pursuit system still responds to target speed exactly as it should, indicating that pursuit is still engaged even when dot coherence is low.

**Figure 5.**
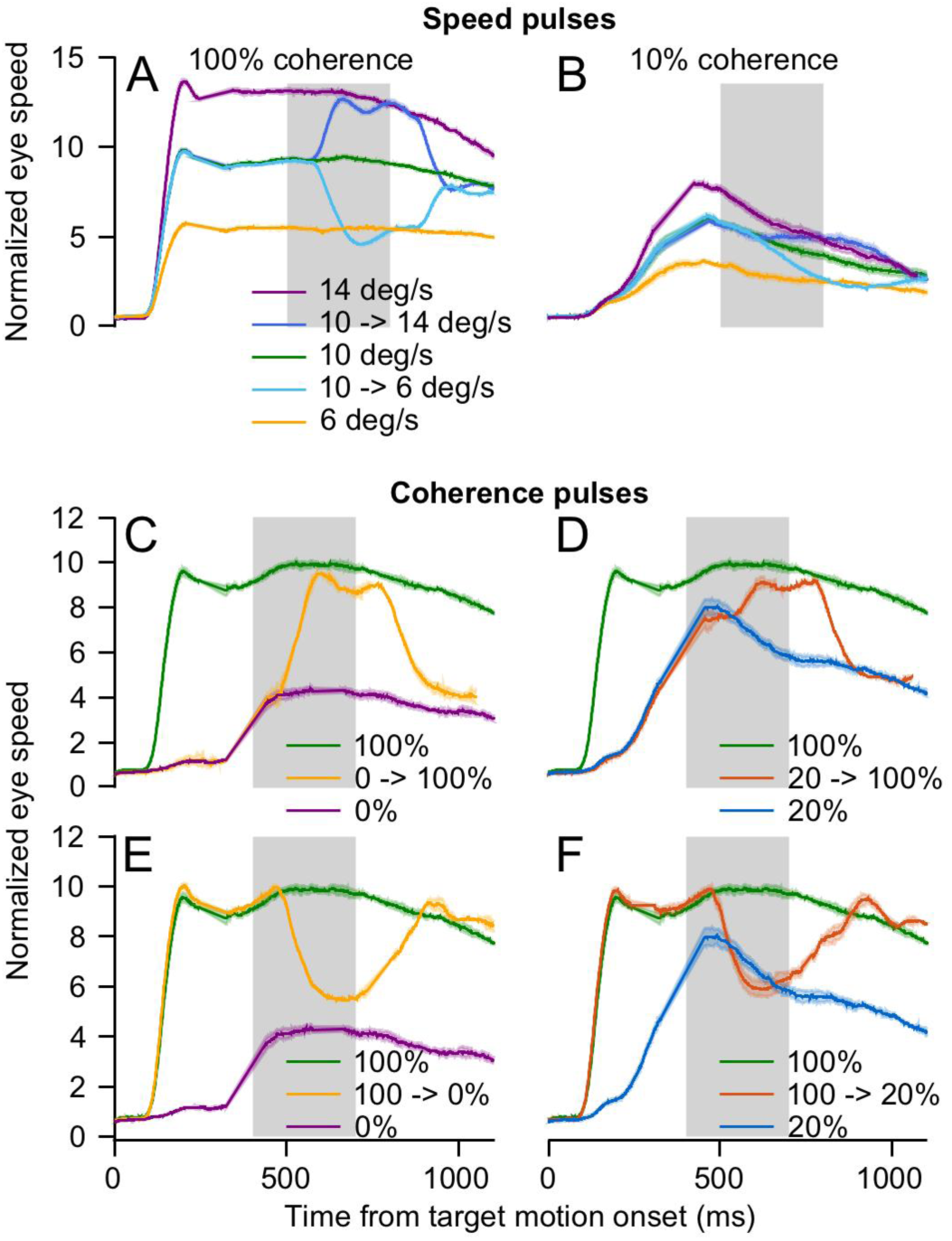
Example eye speed traces for speed and coherence pulses from different starting conditions. Gray shading indicates the duration of the pulses. **A, B**: Speed pulses that increased or decreased target speed from starting conditions of target motion at 10 deg/s and motion coherence of 100% (**A**) or 10% (**B**). In each panel, two of the traces show average eye speeds for speed pulses from 10 to 14 or 6 deg/s and that other three traces show average eye speeds for sustained target motion at 6, 10, or 14 deg/s. **C-F**: Coherence pulses that either increased coherence (**C, D**) from 0% or 20% to 100%, or decreased coherence (**E, F**) from 100% to 20% or 0%. Different colors indicate different starting conditions and pulses. In each panel of **C-F**, one trace shows the average eye speed for a given coherence pulse and the other two traces show the average eye speeds for unperturbed target motions with the two coherences used in that panel. Shaded ribbons around the traces represent standard error of the mean across n=7 to 14 experiments for speed pulses and n=2 to 4 experiments for coherence pulses.

When we introduced changes in target dot coherence, or coherence pulses, during steady-state pursuit by either increasing or decreasing target coherence for 300 ms (Figures 5C-F), pursuit again behaved in a machine-like way. For pulses that increased coherence to 100%, eye speeds converged towards eye speeds during trials that used 100% coherence throughout (Figures 5C, D). For pulses that decreased coherence to 20% or 0%, eye speeds converged towards eye speeds during trials that used the lower coherence values throughout (Figures 5E, F). The responses to coherence and speed pulses indicate that the pursuit system is still engaged and responsive during steady-state tracking, even under conditions of poor tracking due to degraded target motion.

Armed with confidence that the pursuit system is operating like a machine, we can measure directly how low dot coherences perturb the sensory sensitivity of the pursuit system by presenting speed pulses of a range of amplitudes during steady-state tracking. We increased and decreased target speed by increments of 1, 2, 4, and 8 degrees/s for 300 ms starting at 500 ms from motion onset (Figure 6A, pulses indicated by gray shading). In the example eye traces for sets of speed pulses at 100% and 20% coherence (Figures 6B, C), the eye speed responses to the speed pulses are considerably weaker, but still present, when the dot coherence is lower.

**Figure 6.**
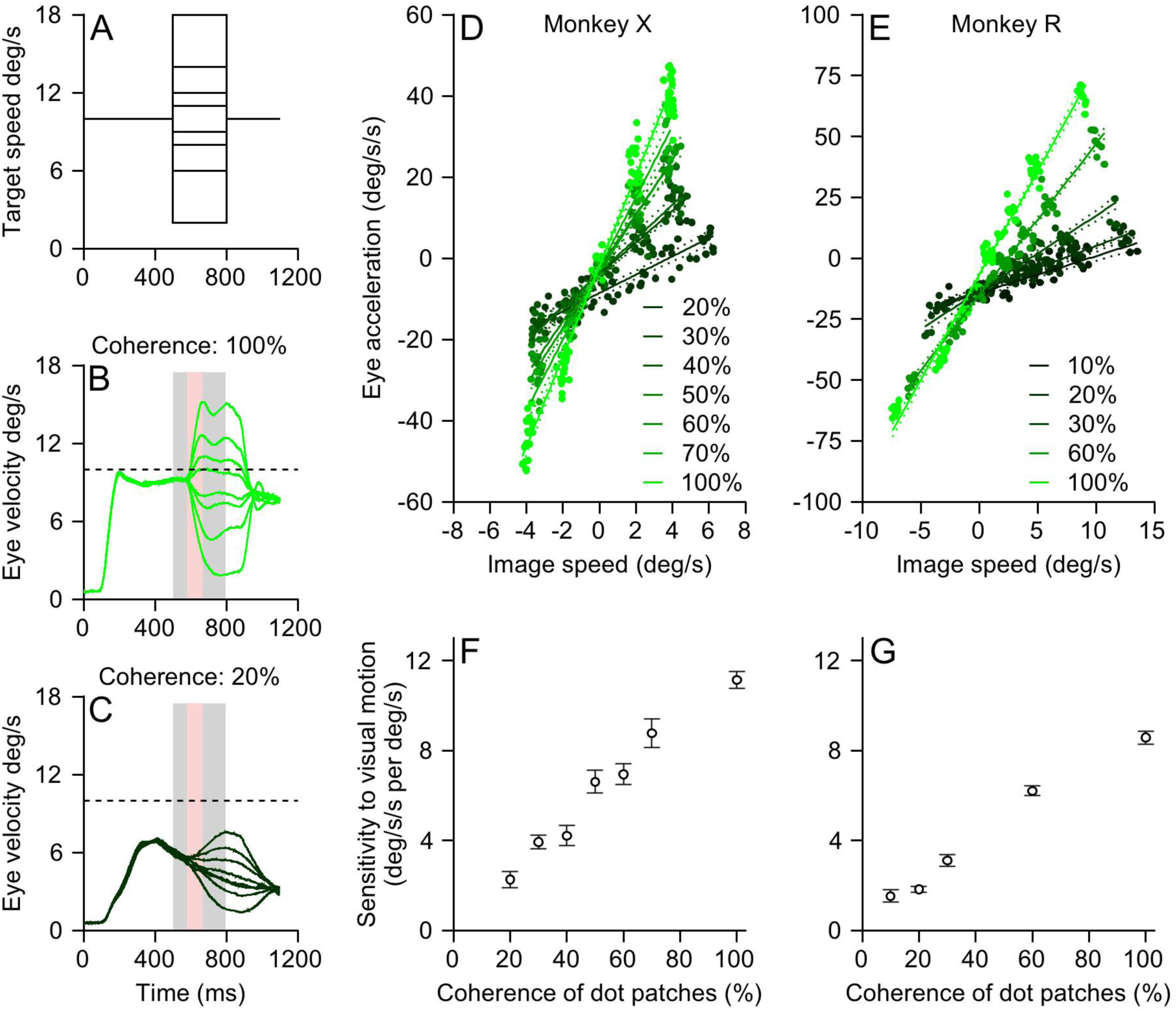
Sensitivity to image motion during deficient steady-state tracking induced by reduced dot coherence. **A**: Target speed trajectories for presentation of speed pulses. **B, C**: Average eye speeds for target speed pulses during steady-state pursuit with target coherences of 100% and 20%, respectively. The grey shading highlights the time of the speed pulse and the pink shading highlights the first 75 ms of the eye response used for analysis. **D, E**: The first 75 ms of eye acceleration plotted as a function of the image speed prior to the beginning of the eye response for two monkeys. The lines show regression fits and the dotted lines represent 95% confidence intervals. **F, G:** The slope of each fit plotted as a function of dot coherence. Error bars represent standard error of the mean across n=4 to 26 experiments in Monkey X and n=3 to 28 experiments in Monkey R. We presented speed pulses during pursuit with 20, 30, 40, 50, 60, 70, and 100% dot coherence for monkey X and 10, 20, 30, 60, and 100% dot coherence for monkey R.

We estimated the sensitivity to image speed at each patch coherence by plotting eye acceleration in the first 75 ms of the response to the speed pulses (interval indicated by pink shading in Figures 6B and C) as a function of the actual image speed in the first 75 ms after the onset of the speed pulse. For both monkeys, eye acceleration was linearly related to image speed at each dot coherence, but the slope of the relationship declined steadily with decreases in dot coherence (Figures 6D and E). Regression analysis of the relationship between eye acceleration and image velocity for each dot coherence shows that the slope of the relationship decreases as a function of dot coherence (Figures 6F and G). It is noteworthy that sensitivity is weak but non-zero even during poor steady-state tracking of targets with low dot coherences. The continued responsiveness of the system to image motion supports the idea that the sensory pathway is still active during poor steady-state pursuit, but eye speed is being pulled down by some other mechanism.

To ask whether changes in dot coherence modulate the motor pathway independently of their effect on the visual representation of motion, we presented changes in dot coherence during steady-state pursuit. At 400 ms after motion onset with 100% coherence dots, we reduced the dot coherence for the remainder of the trial (Figure 7A). The first 75 ms of the eye speed response to the coherence change is the motor system’s response to the reduced coherence in an interval when the image motion driving the current eye movement was consistently near zero no matter the final value of coherence. Example traces in Figure 7B show that decreases in dot coherence caused a decrease in eye velocity, even though the eye had achieved accurate pursuit, with larger drops for lower final values of coherence. We ascribe the decrease in eye speed to modulation of signals in the motor pathways because image motion was always close to zero in the interval just before the decrease in eye speed. For each dot coherence, the decrease in eye velocity was followed by a rebound that we attribute to the visual motion created by the initial deceleration, and finally by a trend towards a new steady-state.

**Figure 7.**
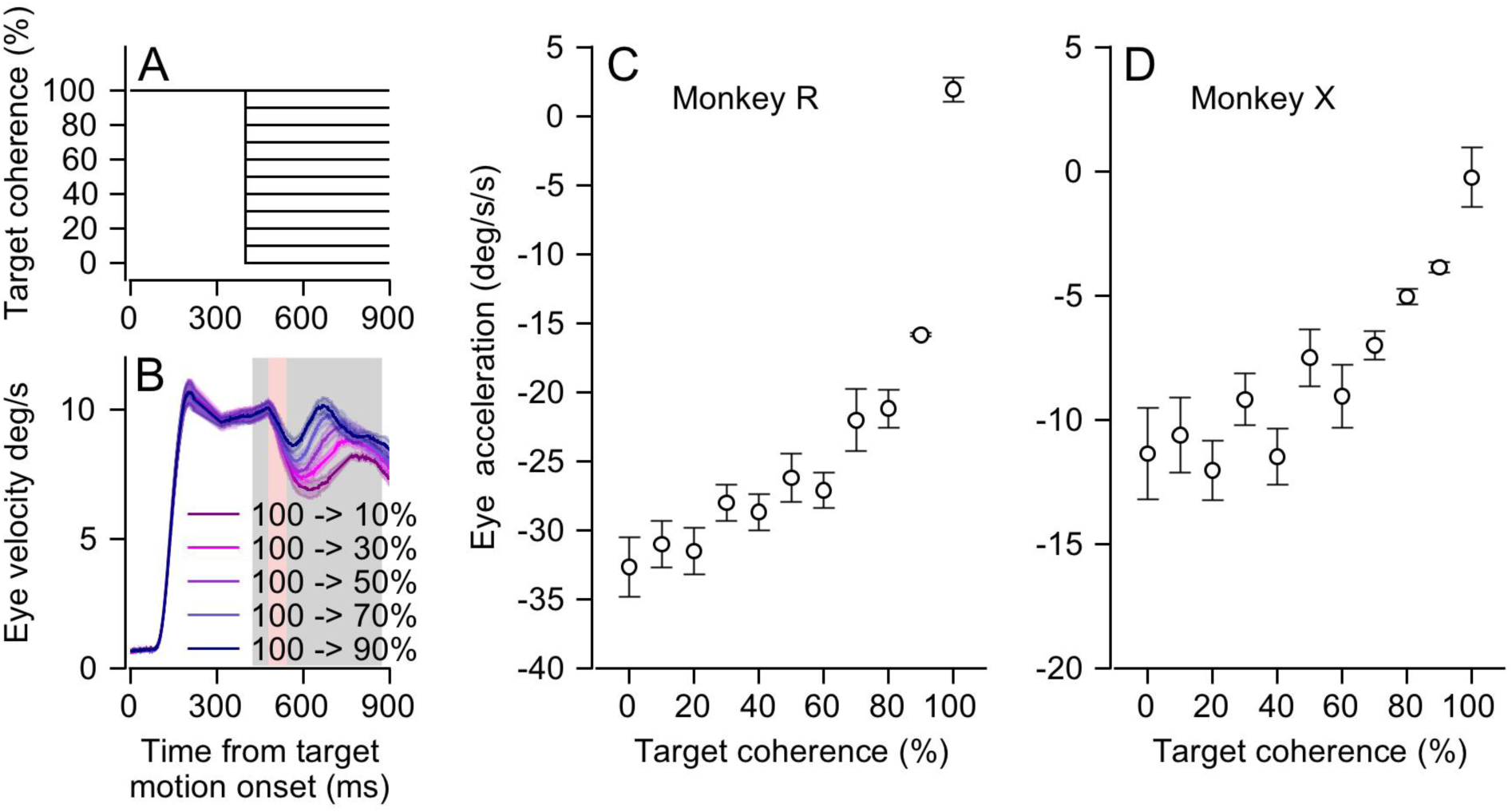
Effect of changes in motion coherence during steady-state tracking. **A**: Trajectory of target coherences. **B:** Eye speed responses to changes in coherence for one monkey. The grey shading indicates the interval when the target coherence was reduced and the pink shading shows the first 75 ms of the eye response used for analysis. The lighter ribbon around each trace represents the standard error of the mean for n=6 experiments for each condition. **C, D**: Average eye accelerations from the first 75 ms of the eye response plotted as a function of target coherence. Error bars are standard errors of the mean across n=6 to 10 experiments for Monkey R and n=8 to 14 experiments for Monkey X.

Steady-state eye velocity eroded in proportion to decreases in dot coherence. We quantified the effect of dot coherence independent of any changes in visual motor drive by plotting the magnitude of the first 75 ms (interval indicated by pink shading in Figure 7B) of eye deceleration as a function of the dot coherence during the reduced coherence (Figures 7C, D). Monkey R showed a steeper effect but both monkeys showed nonlinear increases in the amount of eye deceleration as dot coherence decreased. The responses to coherence pulses are consistent with the idea of a separate modulation for motor pathways and for sensory pathways, and conflict with the possibility that a single change of sensitivity to visual motion accounts for all the effects of dot coherence on pursuit eye movements.

### A model with separate modulation of pursuit initiation and steady-state

Our experiments demonstrated quantitatively different effects of dot coherence on the initiation versus steady-state of pursuit eye movements. The difference in the effect on the two phases of pursuit suggests separate modulation of the neural mechanisms for the two phases. To test this conclusion quantitatively, we have introduced two sites of gain modulation into an existing computational model of pursuit eye movements.

We developed a generative model rather than attempting to fit our data because our goal was a test of plausibility of different model structures. We elected to modify the model of Churchland and Lisberger (2001) because it used two separate mechanisms to produce (1) appropriate dynamical trajectories of eye velocity at the initiation of pursuit and (2) essentially perfect steady-state tracking. In the schematic in Figure 8, the three parallel pathways on the left comprise the sensory pathway and operate on different derivatives of image motion, taken as the difference between the target motion and the eye motion with a lag of 80 ms. The three sensory pathways converge to generate a visual command for eye acceleration that passes through one site of gain modulation denoted by g_1_. The scaled command for eye acceleration is processed by the motor system, represented in the schematic in Figure 8 as a tight loop based on positive feedback of a copy of the final command for eye velocity. The positive feedback loop serves as a mathematical integrator and converts the visual eye acceleration signal into a command for eye velocity. We introduced a second gain term (g_2_) into the positive motor feedback to adjust the quality of steady-state tracking. If the gain of the positive feedback loop is one, then the feedback loop maintains perfect steady-state tracking even in the absence of retinal image motion. If only separate adjustment of both g_1_ and g_2_ can recreate the different effects of dot coherence on the two phases of pursuit, then it would support the idea that there are separate mechanisms for gain modulation, and separate effects of dot coherence, in the visual and motor parts of the pursuit system.

**Figure 8.**
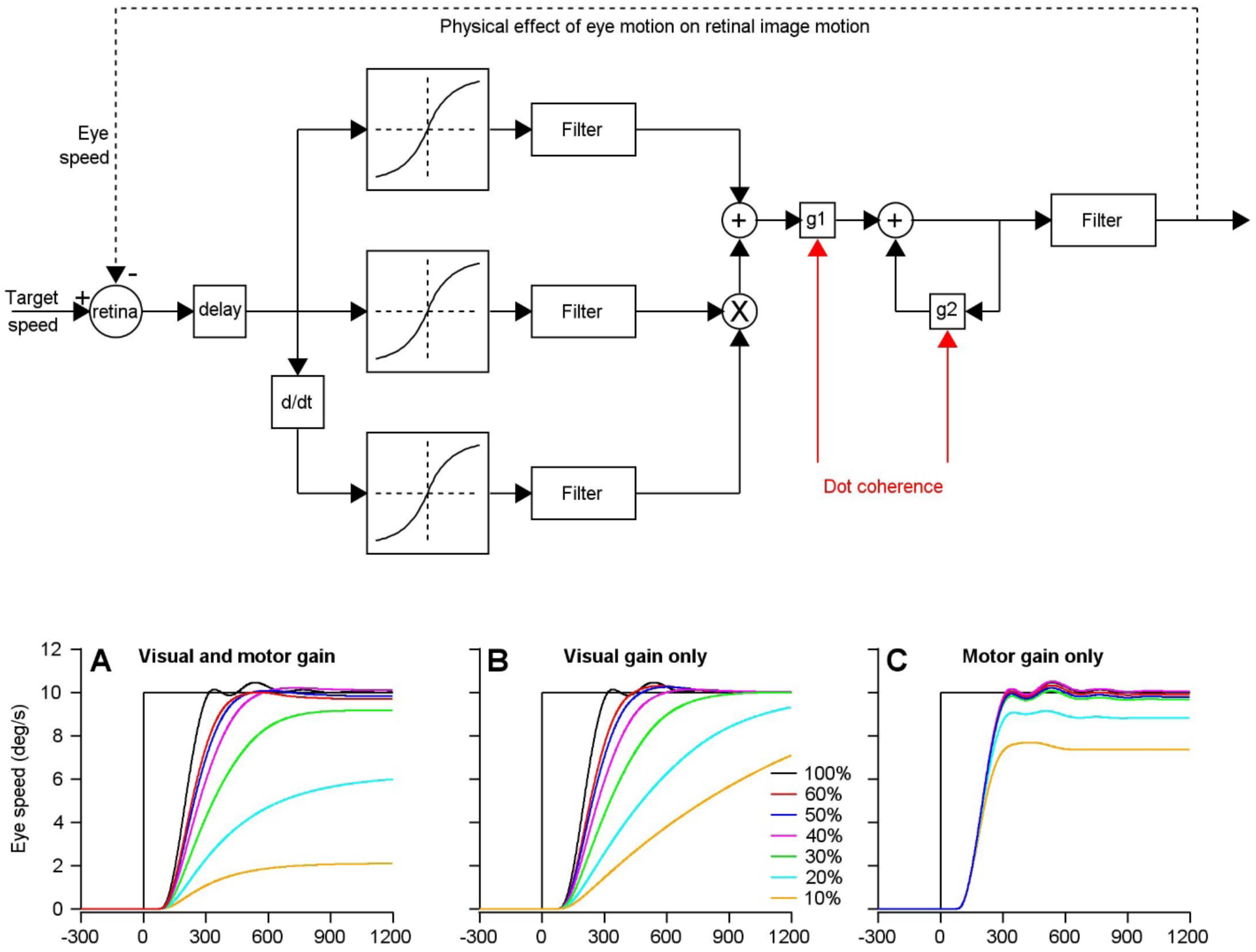
Predictions of a computational model of pursuit behavior showing the effects of changes in gain at two different sites. The schematic at the top shows the model and the red arrows indicate the two sites of gain modulation by dot coherence. The model terms g1 and g2 controlled the gain of visual-motor transmission and the gain of eye velocity positive feedback, respectively. **A-C**: Simulated eye speeds to pursuit of targets with high to low coherence, where different colored traces indicate simulations based on data from different values of motion coherence. The values of g1 and g2 were changed either together (**A**) or individually (**B, C**).

We used the data for monkey X from Figure 3B to define the values of g_1_ and g_2_ as a function of dot coherence. By adjusting these values separately versus jointly, we created three models with modulation of both gains (Figure 8A), modulation only of the gain of visual-motor transmission (Figure 8B), and modulation only of the gain of eye velocity positive feedback (Figure 8C). When we changed the value of g_1_ only, we saw deficits in pursuit initiation but steady-state eye speed gradually increased throughout the target motion and eventually reached target speed when we ran longer simulations (Figure 8B). When we modulated the value of g_2_ only, we observed the expected deficits in steady-state tracking but essentially unaltered initiation of pursuit (Figure 8C). When we modulated both g_1_ and g_2,_ the simulations roughly recapitulate our data (Figure 8A). We conclude that our model of pursuit requires separate modulation of the sensory and motor streams to reproduce the relationships we observed in our data, supporting a conclusion of two sites of gain modulation in the brain.

To perform our computational analysis, we used a phenomenological model that can roughly map onto anatomical systems within the brain. We know that there are more biologically constrained modeling strategies for pursuit that can also output biomimetic behavior. However, we need to acquire biological data about the responses of neurons in MT and the floccular complex as a function of dot coherence before we can evolve our model to be more biological.

## Discussion

We have demonstrated that pursuit eye movements show machine-like effects of altering the coherence within a patch of dots that is used as a pursuit target. The initiation of pursuit and steady-state tracking both are affected by dot coherence. The initiation of pursuit shows decreases in eye acceleration as dot coherence is reduced, while steady-state tracking reaches asymptotic values of eye speed that fall increasingly below target speed as dot coherence is reduced. Several lines of evidence suggest that different mechanisms explain the different effects on the two phases of pursuit. (1) The effects on initiation versus steady-state evolve differently as a function of dot coherence. (2) Sensitivity to image motion is reduced but non-zero during the poor steady-state tracking associated with low values of dot coherence, implying that reduced dot coherence alters the efficacy of a non-visual mechanism that normally operates during steady-state tracking. (3) The effects of dot coherence can be explained by a reliability-weighted competition between sensory evidence and priors based on experience for pursuit initiation (Darlington et al., 2018), but not for steady-state tracking. (4) An accepted computational model of pursuit (Churchland and Lisberger, 2001) can recapitulate our data only if it includes separate modulation of visual mechanisms that produce pursuit initiation versus motor mechanisms that maintain steady-state tracking.

Existing concepts about the organization of the neural circuit for pursuit suggest that visual motion signals from extrastriate area MT are decoded to estimate target speed and provide a command for eye acceleration at the initiation of pursuit (Newsome et al., 1985; Lisberger, 2010). Here, we construct a population response by estimating the response of each of many MT neurons for a given target motion and then plot those responses as a function of each neuron’s preferred speed. Previous papers have shown that both the amplitude of the population response and the preferred speed at the peak of the population response in MT contribute to decoding to estimate target speed (Krekelberg et al., 2006; Darlington et al., 2018). As target speed increases, the peak of the population response shifts towards higher preferred speeds and eye acceleration at the initiation of pursuit increases. However, as the contrast of the moving target is decreased at a fixed target speed, the eye acceleration at the initiation of pursuit decreases (Lisberger and Westbrook, 1985; Yang et al., 2012; Darlington et al., 2017), even though the peak of the population response shifts toward higher preferred speeds while the amplitude of the population response decreases (Maunsell and Van Essen, 1983; Priebe et al., 2003; Priebe and Lisberger, 2004; Krekelberg et al., 2006). We anticipate that appropriately designed recordings from MT will reveal that decreases in dot coherence at a given target speed reduce the amplitude of the population response in MT, possibly without causing a shift in its peak. Thus, the effects of dot coherence on the initiation of pursuit are likely mediated through the same mechanisms as the effects of contrast, and the amplitude of the population response is likely to be the main contributor to reduced eye acceleration for reduced motion coherence.

Our analysis of the effects of context and target contrast on the initiation of pursuit (Darlington et al., 2017, 2018) suggests that two neural signals contribute to the eye acceleration at the initiation of pursuit: (1) a decoded estimate of target speed from MT and (2) a signal from the smooth eye movement region of the frontal eye fields (FEF_SEM_) that controls the gain of visual-motor transmission for pursuit (Tanaka and Lisberger, 2001; Nuding et al., 2009). To account for the effects of context and target contrast on responses in FEF_SEM_, we proposed that the amplitude of the MT population response signals the reliability of the motion and that FEF_SEM_ transforms reliability into visual-motor gain (Darlington et al., 2018). When the contrast of the target is low, a lower amplitude MT population provides a less reliable signal, the output from FEF_SEM_ is smaller, the gain of visual-motor transmission is reduced, and eye acceleration is lower at the initiation of pursuit.

We suggest that the same neural mechanisms operate for dot coherence. We predict that lower values of coherence in a moving dot patch will provide a less reliable motion signal, represented by a smaller amplitude population response in MT, and will lead to lower values of visual-motor gain and weaker eye acceleration in the initiation of pursuit. In this scenario, target speed is estimated as the preferred speed at the peak of the MT population response, neural processing in FEF_SEM_ causes visual-motor gain to be responsive to the amplitude of the MT population response, and the two combine to create a “decoder” that is sensitive to both the peak and amplitude of the population response, as originally suggested by Krekelberg et al. (2006). Importantly, the “decoder” comprises the entire set of pursuit circuits that are downstream from MT. The importance of gain control from FEF_SEM_ in the decoder provided by the full pursuit circuit raises the possibility that gain control could contribute variation in pursuit initiation that arises downstream from the sensory representation of motion, as suggested by Bakhtiari and Pack (2019).

Existing concepts suggest that the neural mechanisms of steady-state pursuit are fundamentally different from those of pursuit initiation. Because steady-state pursuit can be essentially perfect, retinal image motion is eliminated and visual motion signals alone cannot maintain steady-state tracking (Yasui and Young, 1975; Robinson et al., 1986). Instead, considerable evidence supports the conclusion that extra-retinal signals control steady-state tracking. Positive feedback of corollary discharge related to eye velocity through the cerebellar floccular complex seems to be a major contributor to steady-state tracking (Lisberger and Fuchs, 1978; Stone and Lisberger, 1990), explaining how eye velocity can persist if the target is stabilized on the retina during steady-state tracking (Morris and Lisberger, 1987; Lisberger et al., 1987). The failure of steady-state eye speed to match target speed when dot coherence is reduced suggests an effect on the gain of the positive feedback pathways. Two lines of evidence support this conclusion: (1) the pursuit system remains sensitive to visual motion during steady-state tracking of low dot coherences, implying that visual motion should be able to bring eye speed up to target speed, an expectation that our computational model confirms; (2) a step reduction in dot coherence during essentially perfect steady-state tracking causes a decrease in steady-state eye speed even though image motion is unchanged. We conclude that decreases in motion reliability due to low dot coherence alter the gain of the positive feedback of corollary discharge through floccular Purkinje cells. To maintain pursuit at all, the residual sensitivity to visual motion must compete with a tendency for steady-state eye speed to decline, leading to a stable asymptotic eye speed that is lower than target speed. Perhaps the same gain-control mechanism contributes to the results of Deravet et al. (2018), where context had a larger effect on the eye speed at the transition to steady-state pursuit than on the initiation of pursuit. Previous lesion studies in MST (Komatsu and Wurtz, 1988) and FEF_SEM_ (Gottlieb et al., 1994; Tian and Lynch, 1996) have revealed the same conundrum – deficits in steady-state tracking even though visual feedback seems to be intact – suggesting that these areas might have a role in controlling the gain of motor mechanisms as well as of visual-motor transmission.

In conclusion, we developed a behavioral paradigm for investigating the mechanisms behind both the initiation of pursuit and the maintenance of steady-state smooth pursuit tracking. Our data support the hypothesis that sensory and motor pathways driving pursuit are under separate modulated control. Future recordings in area MT and the flocculus of the cerebellum will provide avenues to investigate the mechanisms involved both phases of pursuit as well as the transition from one to the other.

